# Gemini calling! First occurrence of successful twinning in wild, endangered lion-tailed macaques *Macaca silenus* in the Anamalai hills of the Western Ghats, India

**DOI:** 10.1101/2022.08.03.502728

**Authors:** Ashni Kumar Dhawale, Anindya Sinha

**Author notes:** These authors contributed equally to this work.

## Abstract

We recorded the first occurrence of surviving twins in lion-tailed macaques *Macaca silenus* from the Anamalai hills of the Western Ghats, India. The Puthuthottam population of lion-tailed macaques has historically been restricted to a rainforest fragment measuring 92ha, situated adjacent to human settlements, however, direct interactions between macaques and humans have been observed only in the last ten years. The population now visit settlements at a rate of 0.52/ day, and exploits anthropogenic foods. We recorded two sets of twins from the Puthuthottam population within a six-month period. Two previous cases of twins have been observed in this population since 2000, however, in both cases the twins did not survive beyond a few weeks. We followed and opportunistically collected ad libitum behavioural data on both sets of twins for a year between March 2019 to March 2020. Both mothers carrying the twins used the ground substrate extensively, however the mother with the younger set of twins also used the canopy and other precarious substrates such as cable wires. Our report shows that twinning occurs in lion-tailed macaques and twins are able to survive successfully. This report also supports previous evidence that twinning may occur in higher frequencies as a direct response to provisioning, with the mothers’ behaviour contributing to the successful survival of both twins.

## Introduction

Both humans, and non-human primates have, through evolutionary history, exhibited the ability to undergo *‘twinning’*, or “the gestation of multiple embryos [at the same time]” (Varella, 2019). While generally, primates give birth to a single offspring, records show the occurrence of twinning in many species across primates, including Chimpanzees *Pan troglodytes* and *Pan troglodytes schweinfurthii* (Ely et al., 2006; Matsumoto-Oda, 1995; Peacock & Rogers, 1959), gorillas *Gorilla gorilla* (Rosen, 1972) with a case of conjoined twins (Langer et al., 2014), orangutans (Goossens et al., 2011; Hudson, 1976), vervet monkeys (Pollack & Raleigh, 1994), langurs *Presbytis melalophos* (Bennett, 1988) and *Presbytis entellus* (Winkler et al., 1989), capuchins *Cebus apella* (Leighty et al., 2003) and *Cebus capucinus* (Manson & Perry, 2000), marmosets (Haig, 1999), pygmy loris *Nycticebus pygmaeus* (Jurke et al., 1998), lemurs (Morland, 1990; Parga & Nansen, 2019) and galagos *Galago crassicaudatus umbrosus* (Pasztor & Van Horn, 1976; Tartabini, 1991). Twinning is the norm in callitrichine primates, having evolved through genetic changes resulting in a propensity for multiple pregnancies (Harris et al., 2014), however, it is incredibly rare in Great Apes and Old World monkeys. Studies on certain species of macaques, including the Japanese macaque *Macaca fuscata* (Sugiyama et al., 2011), rhesus macaque *Macaca mulatta* (Bercovitch et al., 2002) and Tibetan macaque *Macaca thibetana* (Xia et al., 2012) have been observed twins, however, it is noteworthy that these reports are specifically from artificially provisioned populations, and even so, twinning rate is estimated at 0.02-1%. In an entirely anomalous case, a population of Formosan macaque *Macaca cyclopis* in Taiwan exhibits a high prevalence of polythelia or the presence of supernumerary nipples, which is observed to be linked to the high incidence of twinning (Hsu et al., 2000). In cases of twinning in catarrhine primates, more often than not, twins are not observed to survive long, or only one in the set survives, especially in the wild.

In this paper we present the first record of surviving twins in the lion-tailed macaque, from the Anamalai Hills in the Western Ghats of India. The lion-tailed macaque is an endangered primate endemic to the Western Ghats of southern India. Currently, the total population is estimated at 3000-3500 individuals (Singh et al., 2009) and occurs, across its entire distribution, scattered in pockets of wet evergreen rainforest. Known as the only truly arboreal macaque species, the lion-tailed macaque is a habitat specialist, being highly adapted to its rainforest habitat. While primarily frugivorous, their diet also includes invertebrates and on some rare occasions, small vertebrates. The lion-tailed macaque has a unique life-history, with females reaching sexual maturity at a relatively late age, having a long inter-birth interval of three years, and carrying their infant for nearly a year (Erinjery et al., 2017; Kumar, 2013).

For the species, twinning has been reported in only one instance from an 81-year record of captive lion-tailed macaque individuals in North American zoos (Lindburg et al., 1989). Additionally, two previous cases of twins have been observed in the Anamalai Hills population (Kumar pers. Obs.), however, in both cases the twins did not survive beyond a couple of weeks. We now present observations on not one, but two sets of surviving twins born within a six-month period of each other, in a unique lion-tailed macaque population.

## Methods

### Field Site

Our study was conducted in the Valparai plateau 10° 19’ 39.22”N, 76° 57’ 18.98”E, situated in the Anamalai hill range of the Western Ghats, in the southern Indian state of Tamil Nadu. The 220 km^2^ Valparai plateau is mainly a plantation landscape, dominated by tea and coffee, with remnant wet evergreen rainforest fragments measuring between <11ha to >200ha (Mudappa & Shankar Raman, 2007; Muthuramkumar et al., 2006). These forest fragments, despite being surrounded by a human-dominated habitat matrix, harbour many wild species including the Asian elephant, leopard, sloth bear, mouse deer, a variety of small carnivores, invertebrates and birds, and the endemic lion-tailed macaque.

### Study Subjects

One particular forest fragment, the Puthuthottam forest fragment, retains a population of ∼190 lion-tailed macaques divided into five troops. The Puthuthottam macaques are functionally restricted to this 92ha forest fragment, which is surrounded by tea plantation and neighbours the town of Valparai. Over the last three decades, the population has steadily increased (Singh et al., 2001, 2002; Umapathy, 1998; Umapathy & Kumar, 2003) and more recently, all troops have begun to explore the surrounding human-dominated landscape. This expansion into human-habitation has allowed the macaques to utilise anthropogenic foods primarily through home- and garbage raiding (Dhawale et al., 2020). Efforts made by a local NGO, Nature Conservation Foundation (NCF), have kept direct provisioning by tourists to a minimum.

### Field Methods

Each of the five troops, coded as BT, HAN, PAP, RT and NTT, were systematically followed at per-determined sampling intervals between March 2019-March 2020 (15±5 days/month). Each time we observed a set of twins, ad libitum behavioural data was collected on the individual and her associations with the infants and other troop members. Since the BT troop contained ∼90 members, we were unable to individually identify all of them, however, members of all other troops were individually identified using distinctive markers, such as facial markings, visible injuries, or other visible abnormalities including, missing body parts or swellings. Our observations on each set of twins are described below.

## Observations

### BT Twins

We first sighted a pair of twins (Figure 1) on March 22, 2019, in the BT troop, being carried by an adult female belonging to an older age class. The adult female, henceforth TM, was first spotted on the ground substrate, foraging for insects among the leaf-litter. Both infants were alert and seemed to be about one to two weeks of age. During the first two months of observation between April-May 2019, one infant held on ventrally to the mother while the second infant reached over the first infant’s head to hold onto the mother’s opposite side (Figure 1, right panel), each infant positioned slightly away from the mother’s midline. The infants did not release hold of TM during this initial period, thus, determining the sex of the infants was not possible. Both infants showed oral rooting and suckled from both nipples; as we could not identify the infants, we could not determine preference.

**Fig 1.**
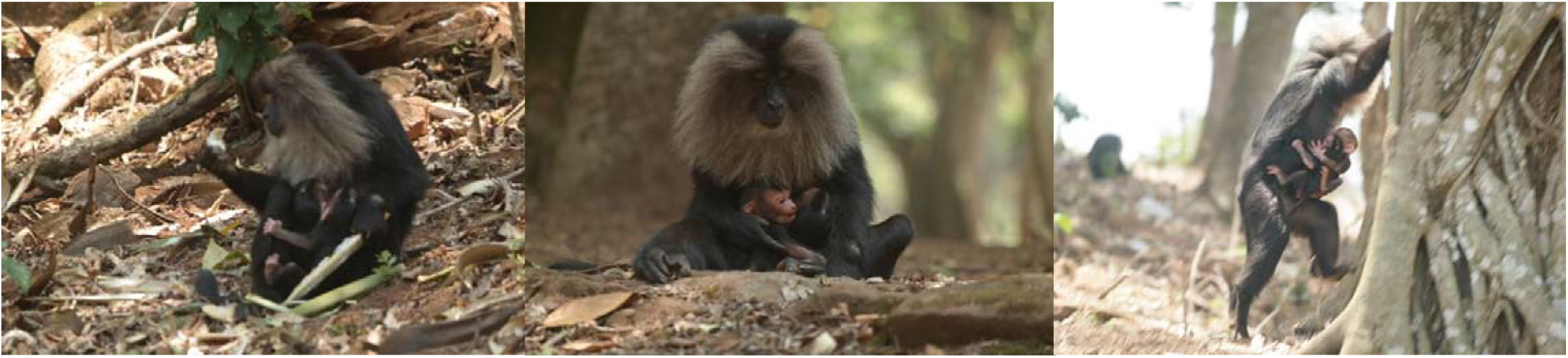
BT Twins first recorded in March 22, 2019, with mother TM

TM moved around in a quadrupedal manner, without providing support to the infants, however, she was observed to mostly utilise the ground substrate or the lower canopy (within 5m of the ground). She also associated with other members of the troop very rarely, preferring to move away if an individual approached. When she was first spotted, TM was also nursing a wound on her left wrist, presumably from unrelated causes. Due to the heavy monsoon season in the Western Ghats, between June-August 2019, we could not conduct detailed behavioural observations on TM except for intermittent monitoring to ensure survival of the twins.

The twins had grown substantially by September 2019, and TM seemed to be visibly overburdened by the weight of the infants. She was now observed to walk with one hand supporting the twins’ heads and rarely left the ground substrate. The infants were more active and would release hold of the mother, however, they stayed within 1m of TM at all times. At this time, we were able to visually determine the sex on the infants, one male and one female.

The twins were observed to play among themselves, even while being held, and perform facial gestures at one another and at TM, who would spend most of her time grooming them, feeding, resting and moving only on the ground or very low in the vegetation. She was also seen to interact more often with other adult females and even juveniles in the troop, offering grooming and being groomed by the others. Towards the end of the year, in March 2020, the twins could only be identified when they sat close with TM during resting bouts. At other times, they were observed to join in play sessions consisting of up to 30 juveniles, where individual identification was impossible.

Lion-tailed macaques residing in Puthuthottam require to every so often cross the main road which cuts through the fragment. We have observed adult females ventrally carrying juveniles up to three years of age, when crossing such large open spaces (Dhawale pers. Obs.), especially when using ‘canopy bridges’ or aerial bridges affixed on the upper canopy of trees spanning the road, a conservation measure set in place by NCF. During one instance of such a crossing, we observed TM having difficulty transporting the twins across the bridge causing her to stay back as the rest of the troop crossed; eventually, an older adult female approached TM, exchanged affiliative behaviours including lip smacking and touch, and carried one infant followed closely by TM carrying the other. Once they had crossed the bridge, TM was seen to retrieve the second twin and move for a short distance before resting, while the twins moved away to play with members of their cohort in the branches nearby. As part of a larger behavioural study on the Puthuthottam population of macaques, we intermittently monitored the twins and established their survival until the time the manuscript was written.

### RT Twins

The second set of twins (Figure 2) was sighted on 3 October, 2019 in the RT troop, being carried by adult female SE belonging to a relatively younger age class (not primiparous). The infants were less than a week old and appeared to be smaller in size than a singleton of the same age. In this case as well, both infants were alert and actively looking around, but held by the mother at all times. Similar to the BT twins, one infant held on to the mother directly, while the other reached over the sibling’s head and held onto the mothers opposite side (Figure 2 right panel). The infants were observed to suckle simultaneously from both nipples.

**Fig 2.**
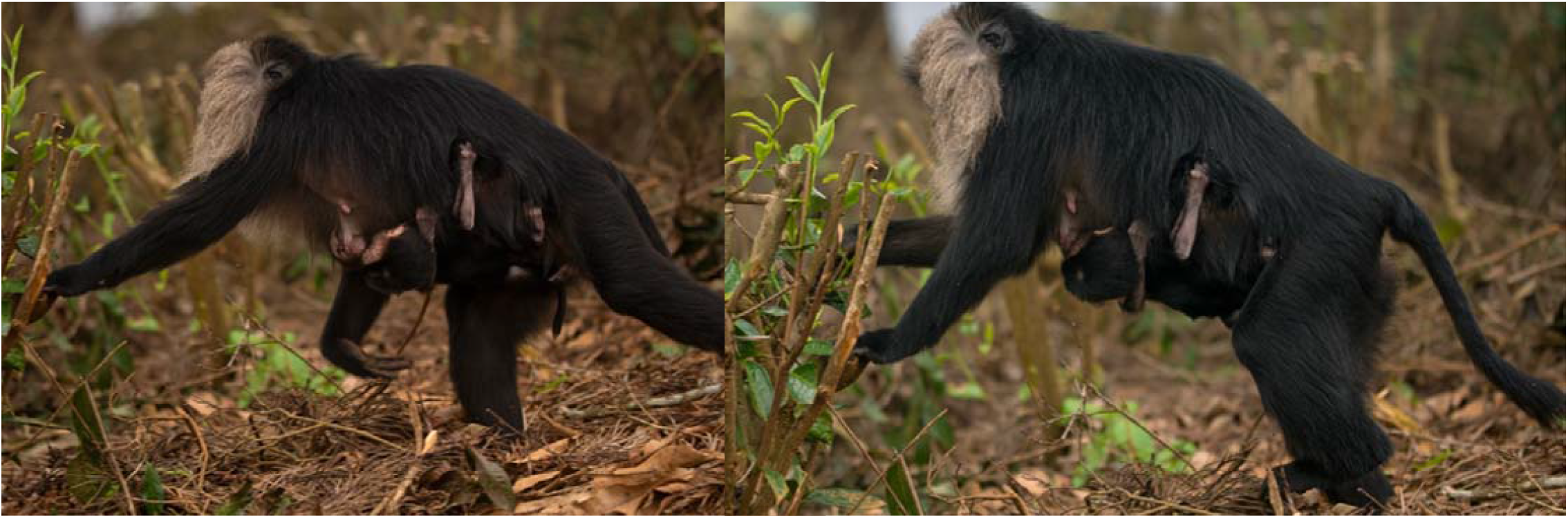
RT Twins first recorded in October 3, 2019, with mother SE

During the initial days of observation, SE was observed to use all substrate types from ground to upper canopy, and even traversed a cable wire (Figure 3), without providing support to the twins, who remained holding on to her ventral side. As time progressed and the twins grew in size and SE was seen to remain on the ground substrate or rooftops of houses, avoiding vegetation and the upper canopy entirely. Both twins were observed to be male.

**Fig 3.**
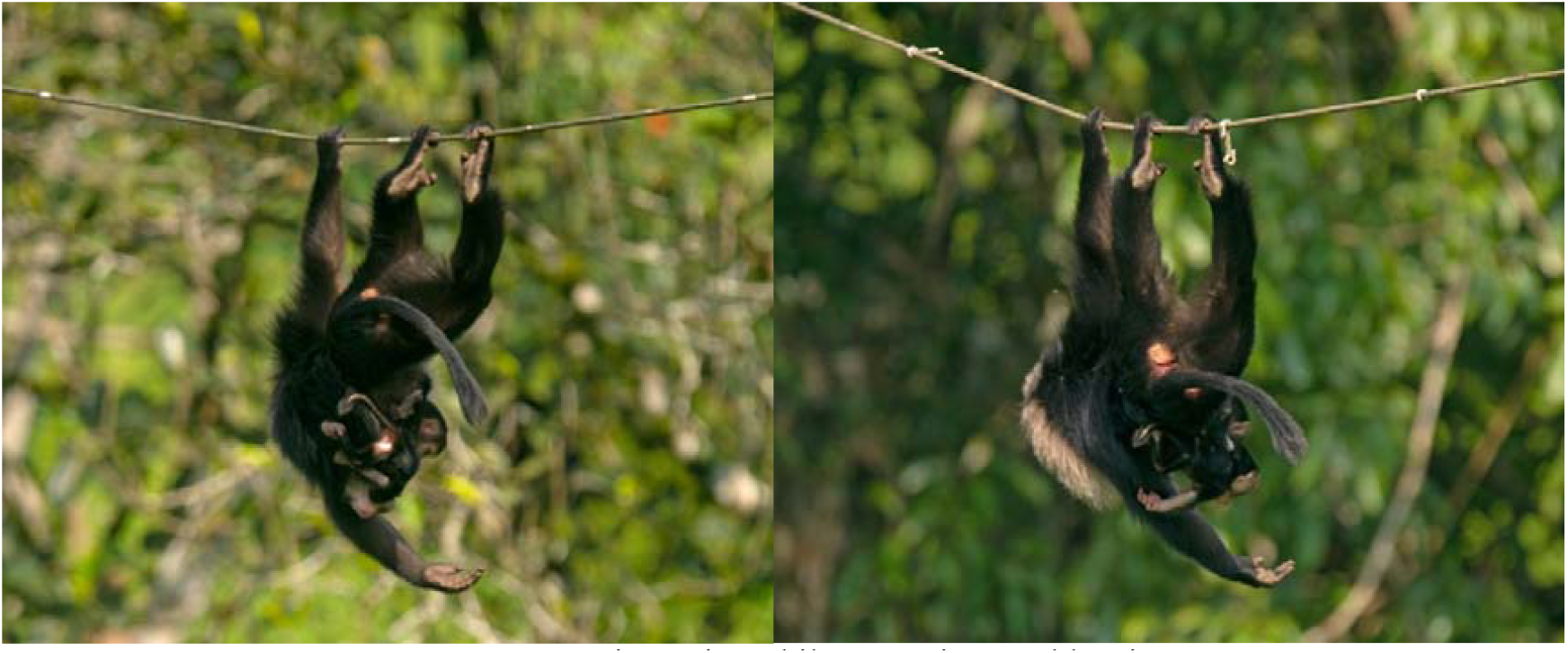
SE carrying twins while traversing a cable wire

Throughout the study period, SE was observed to interact with other members of the troop, same as if she had a single offspring. We continued to intermittently monitor the RT Troop throughout 2020, and recorded the survival of the twins beyond the infant age class.

## Discussion

An analysis of 70 primates revealed that twinning propensity was higher in species having smaller body size with short gestation and weaning periods, a diet comprised primarily of insects which provide non-seasonal access to nutrition (Chapman et al., 1990 as described by Harris et al., 2014) and specific social dynamics like alloparenting (Harris et al., 2014). In larger mammals, environmental conditions such as social organisation has been known to influence birth numbers, and individuals of higher dominance rank are observed to have more success in carrying out multiple pregnancies (Sugiyama et al., 2011). More recently, access to high energy foods through artificial provisioning, whether in captivity or in wild setting, has been repeatedly associated with the increase in twinning frequency (Bercovitch et al., 2002; Peacock & Rogers, 1959; Sugiyama et al., 2011; Xia et al., 2012). Lastly, twinning has also been associated with age (Harris et al., 2014; Peacock & Rogers, 1959) and genetic predisposition (Bennett, 1988; Hsu et al., 2000; Sugiyama et al., 2011)

Apart from its recent utilisation of anthropogenic foods through home- and garbage-raiding, the lion-tailed macaque does not adhere to the ecological criteria under which twinning is observed. In the case of our reported twin sightings, the mothers were from two entirely different troops with nearly non-overlapping home ranges (unpublished data), ruling out the possibility of a genetic basis for the observed pattern. Furthermore, dominance rank, known to influence living conditions such as assess to food and beneficial social interactions, has been linked with increased chance of bearing twins, however, while TM seemed to be a dominant individual, based on ad libitum approach-retreat observations, SE held an intermediate ranking. Finally, while TM was an older adult female, SE belonged to a relatively young age-class, based on nipple length and other morphological features. Thus, we believe the primary driver for twinning in this species was the presence of energy rich, non-seasonal, easily accessible anthropogenic foods.

Anthropogenic foods are present in the landscape in the form of open garbage dumpsters, litter on the ground, and found within homes. Both lion-tailed macaque mothers were observed to forage extensively at these locations, which allowed them to overcome the burden of transporting the twins to higher substrates. Both BT and RT inherently used the ground and rooftop substrate most often, maintaining relatively small home ranges c. 309.1 and 16.7 ha respectively, and remaining within close vicinity of human habitations at all times. Walking short distances, an average of 500m per day, allowed both mothers to locomote easily while supporting the twins with one hand. Beyond 6 months of age, both sets of twins were observed to walk short distance alongside the mother. Additionally, free access to anthropogenic foods and a disproportionately high presence of invertebrates in this population’s diet (Dhawale et al., 2020) perhaps aided the mothers in milk production to compensate for the additional requirements. The differences in dominance ranks across the two individuals perhaps influenced the observed varied frequencies of intra-troop social interactions, with SE, the intermediate female, having to maintain her position or potentially rise in rank through affiliative interactions, to reap the associated benefits.

Therefore, we believe a combination of environmental conditions and unique behavioural adaptations exhibited by the Puthuthottam population allowed for the successful survival of the twins. This report confirms that twinning is possible in the species and under certain environmental conditions, survival is likely.

## Notes

### Competing Interest Statement

The authors have declared no competing interest.

